# Oxidative stress profiles of lymphoblasts from Amyotrophic Lateral Sclerosis patients with or without known SOD1 mutations

**DOI:** 10.1101/2022.03.03.482309

**Authors:** Teresa Cunha-Oliveira, Daniela Franco Silva, Luis Segura, Inês Baldeiras, Ricardo Marques, Tatiana Rosenstock, Paulo J. Oliveira, Filomena S.G. Silva

## Abstract

Amyotrophic lateral sclerosis (ALS) is a fatal and rapidly progressing neurodegenerative disease that affects motor neurons. This disease is associated with oxidative stress especially in mutant superoxide dismutase 1 (mutSOD1) patients. However, less is known for the most prevalent sporadic ALS due to a lack of disease models. Here, we studied oxidative the stress profiles in lymphoblasts from ALS patients with mutSOD1 or unknown (undSOD1) mutations.

mutSOD1 and undSOD1 lymphoblasts, as well as sex/age-matched controls (3/group) were obtained from Coriell and divided in 46 years-old-men (C1), 46 years-old-women (C2) or 26/27 years-old-men (C3) cohorts. Growth curves were performed, and several parameters associated with redox homeostasis were evaluated, including SOD activity and expression, general oxidative stress levels, lipid peroxidation, response to oxidative stimulus, glutathione redox cycle, catalase expression, and activity, and Nrf2 transcripts. Pooled (all cohorts) and paired (intra-cohort) statistical analyses were performed, followed by clustering and principal component analyses (PCA).

Although a high heterogeneity among lymphoblast redox profiles was found between cohorts, clustering analysis based on 7 parameters with high chi-square ranking (total SOD activity, oxidative stress levels, catalase transcripts, SOD1 protein levels, metabolic response to mM concentrations of tert-butyl hydroperoxide, glutathione reductase activity, and Nrf2 transcript levels) provided a perfect separation between samples from healthy controls and ALS (undSOD1 and mutSOD1), also visualized in the PCA analysis.

Our results show distinct redox signatures in lymphoblasts from mutSOD1, undSOD1, and healthy controls that can be used as therapeutic targets for ALS drug development.

**Highlights:**

⍰ Lymphoblasts from ALS patients present altered redox properties
⍰ Redox profiling evidenced total SOD and glutathione reductase activities, SOD1 protein levels, DCF fluorescence, catalase and Nrf2 transcripts, and tert-butyl hydroperoxide cytotoxicity, as discriminant features for experimental groups.
⍰ Lymphoblast redox profiles may be helpful for patient stratification and precision medicine

## 1 Introduction

Amyotrophic lateral sclerosis (ALS - Amyotrophic Lateral Sclerosis), also known as Lou Gehrig’s disease or Charcot disease, is the most common motor neuron disorder with a worldwide incidence of 2 cases per 100,000 per year[1–4]. This late-onset disease is characterized by selective loss of upper and lower motor neurons in the cerebral cortex, brainstem and spinal cord, causing progressive muscle weakness, atrophy and paralysis, leading to death due to respiratory failure[1, 2, 5]. There is no cure for ALS, and most patients survive in average 2-3 years after diagnosis[6, 7]. Only two drugs are currently approved by the US Food and Drug Administration for ALS treatment, including riluzole, a neuroprotective glutamate antagonist that extends the survival rate over 2-3 months, and edaravone, an antioxidant that delays ALS progression only beneficial in a subset of patients with early-stage ALS[8–11].

Familial ALS (fALS) is associated with 5–10% of the cases, while sporadic ALS (sALS), the most predominant form, accounts for the rest. The etiology of sALS remains unknown and may be associated with several risk factors, including age, smoking habits, body mass index, physical fitness, dietary factors (e.g. intake of antioxidants), and occupational and environmental risk factors[12–17]. However, none is considered as directly triggering the development of ALS[12–16]. Therefore, individual susceptibility factors and external exposure to environmental factors can determine the onset of the disease[18, 19]. Over 50 disease-modifying genes have been described in ALS[20], and the superoxide dismutase type-1 (SOD1) mutation (mutSOD1) was the first mutation identified to be related to ALS[21], accounting for 20% of familial cases and 1-7% of sporadic cases[21–28]. Other mutations associated with ALS include C9orf72, TDP43 and FUS[6]. However, since neither mutations nor risk factors alone fully describe the etiopathogenesis of ALS, a gene-time-environment model has emerged, in which the development of ALS is a multistep process, with the genetic background being one of the several triggering factors [18]. The precise pathophysiological mechanism of ALS remains unclear, but since sALS and fALS present similar clinical features, these forms may share at least some pathological mechanisms [2]. Among the molecular mechanisms proposed, glutamate excitotoxicity, dysregulation of transcription, altered RNA metabolism, defective axonal transport, protein misfolding and aggregation, endoplasmic reticulum stress, disrupted protein trafficking, proteasome impairment, inflammation, activation of cell death signals, mitochondrial dysfunction, and oxidative stress are the most discussed[2, 29–32].

Higher levels of oxidative damage to DNA, proteins, and lipids were reported in postmortem neuronal tissue[33–37], cerebrospinal fluid (CSF)[38–41], plasma,[42–44], and urine[45] from ALS patients. Although these results provide evidence of oxidative stress involvement in ALS pathogenesis, it is not clear if oxidative stress represents a primary cause or a secondary consequence of the pathology, and whether it is present early or later in the disease. This type of study in ALS patients has several limitations, including the short life span of the patients, which does not allow following markers of oxidative damage for a long time (longitudinal study), and the impossibility of evaluating oxidative damage markers in early disease, as the initial stage of the disease progresses in a subclinical manner, sometimes years before diagnosis[6].

SOD1 is a predominantly cytosolic Cu-Zn metalloprotein responsible for eliminating superoxide radical. This enzyme plays a key role in regulating oxidative stress, energy metabolism, cellular respiration, and cell damage[46–50]. In addition, some of the environmental ALS risk factors, including exposure to agricultural chemicals, heavy metals, excessive physical exertion, chronic head trauma, and smoking, have been associated with oxidative stress, suggesting that the development of ALS can be facilitated by any factor that favors a prooxidative state[51]. However, the mechanisms leading to oxidative stress in ALS patients without SOD1 mutations must be clarified. Until now, mechanistic insights have been mainly obtained from animal models, such as the mutSOD1 mouse models.

Profiling the oxidant/antioxidant imbalance in ALS patients may be helpful to find fingerprints and therapeutic targets for ALS, ultimately allowing patient stratification. Since ALS is a multisystemic disease, lymphoblasts have been increasingly used for this purpose since they share molecular mechanisms, such as protein aggregation, mitochondrial dysfunction, with degenerated motor neurons[52]. Lymphoblasts allow the analysis of multiple variables from the same biological source, identifying internal biological correlations and testing personalized therapeutic strategies. Here, we characterized oxidative stress profiles in lymphoblasts obtained from ALS patients with characterized SOD1 mutations, no characterized (undetermined) SOD1 mutations, and their respective age-and sex-matched controls to identify redox discriminant markers.

## 2 Materials and Methods

### 2.1 Experimental design

Lymphoblast cell lines, established by Epstein-Barr transformation, from 3 patients with mutSOD1 and 3 patients without identified SOD1 mutations (undSOD1), and 3 controls of the same age and sex were obtained from the cell line repository at Coriell Institute for Medical Research, USA (www.coriell.org). The groups were analyzed in pools, or divided into 3 cohorts, each containing 1 sample from healthy control, 1 sample from one undSOD1 patient and 1 sample from one mutSOD1 patient: **C1**-Females 46 years old; **C2**-Males 46 years old; **C3**-Males 26/27 years old (Supplemental Table 1).

### 2.2 Cell culture and analysis of cell proliferation

Lymphoblasts were grown in RPMI 1640 medium from Sigma-Aldrich (Saint Louis, MO, USA, R6504) supplemented with 2 g/L sodium bicarbonate, 15% (v/v) fetal bovine serum and 0.5% (v/v) of penicillin-streptomycin plus amphotericin B in T25 or T75 flasks, in an upright, at 37°C in a humidified atmosphere of 5% CO_2_. The cells were counted using a Bio-Rad^®^ TC20 Automated Cell Counter (Bio-Rad, Hercules, CA, USA) and kept in culture at a density of 0.4 to 1.2 x10^6^ cells/mL. For cell counting, 10 μL of each cellular suspension was diluted in 0.4% Trypan blue solution from Gibco (Hampton, USA, 15250-061) in a ratio 1:1.

The lymphoblasts were sub-cultured at an initial density of 0.3 x 10^6^ cells/ml in T25 flasks with a final volume of 10 mL and maintained in culture up to 96 h to perform proliferation curves. Viable cells were counted after 24, 48, 72 and 96 h of cell growth, and the doubling time of each cell line was calculated based on the exponential (Malthusian) growth model.

Every 2-3 days, the cultures were diluted and re-fed with fresh medium according to the rate of cell growth and the required number of cells needed for the experiments, and once per week, the medium was entirely renewed after cell centrifugation. Before all experiments, lymphoblasts were plated at 0.4 or 0.7 x 10^6^ cells/mL in RPMI medium and incubated for 48 h or 24 h, respectively, to secure a cell density of around 1 x 10^6^ cell/mL in each assay. Considering that doubling times can affect cell phenotype and responses, cells were discarded after 30 population doublings from the initial culture provided by Coriell, and new cultures were started from aliquots frozen at low cell passage.

### 2.3 Analysis of gene expression

Total RNA was extracted from dry pellets containing ~5 x10^6^ cells, using RNeasy mini kit (Qiagen, Düsseldorf, Germany), following the manufacturer’s protocols, and quantified using a Nanodrop 2000 (Thermo Fisher Scientific, Waltham, MA, USA), confirming that the A260/280 ratio was higher than 1.9. RNA was converted into cDNA using the iScript cDNA synthesis kit (Bio-Rad, Hercules, CA, USA), following the manufacturer’s instructions. RT-PCR was performed using the SsoFast EvaGreen Supermix, in a CFX96 real time-PCR system (Bio-Rad, Hercules, CA, USA), with the primers defined in Supplemental Table 2, at 500 nM. Amplification of 12.5 ng of cDNA was performed with an initial cycle of 30 s at 95.0°C, followed by 40 cycles of 5 s at 95°C plus 5 s at the annealing temperature (Ta) shown in Supplemental Table 2. At the end of each cycle, EvaGreen fluorescence was recorded to determine Cq. After amplification, the melting temperatures of the PCR products were determined by performing melting curves. For each set of primers, amplification efficiency was assessed using ten-fold dilutions of a pool of all samples, and no template and no transcriptase controls were ran. Relative normalized expression was determined by the CFX96 Manager software (v. 3.0; Bio-Rad), using *TBP*, *YWHAZ*, *PUM1* and *B2M* as reference genes.

### 2.4 Determination of total SOD activity

Total SOD activity (considering all SOD isoforms) was determined using a colorimetric kit by Enzo Life Sciences (ADI-900-157), following the manufacturer’s instructions. Briefly, lymphoblasts were seeded at 400,000 for 48 h in T75 flasks containing with 30 mL RPMI medium, to secure a cell density of around 1×10^6^ cells/mL in each assay. After 48 h, cells were harvested, washed with ice-cold 1x PBS, and lysed following the manufacturer’s instructions. Protein concentration was determined using the bicinchoninic acid method (BCA), 4,4’-dicarboxi-2,2’-biquinoline)[53], at 450 nm in a BioTek Cytation 3^™^ microplate reader (BioTek Instruments Inc., USA). Cell lysates containing 25 μg protein were incubated with 150 μL of Master Mix (composed of 10x SOD buffer, WST-1 reagent, xanthine oxidase and H_2_O milliQ), 25 μL of xanthine 1x and the absorbance was read at 450 nm in a BioTek Cytation 3^™^ multiplate reader (BioTek Instruments Inc., USA). Results were expressed as % of the pool of controls.

### 2.5 Western blotting

Total cellular extracts corresponding to 30 x 10^6^ cells were centrifuged at 259 x g, at room temperature, for 5 min, and the resulting pellet was washed once with PBS. The cellular pellet was resuspended in cell lysis buffer containing 20.4 mM ethylenediaminetetraacetic acid (EDTA), 146.9 mM ethylene glycol-bis (2-aminoethylether) N, N,’N, ’N-tetraacetic acid (EGTA), 1 mM calcium glycerophosphate, 1 mM sodium orthovanadate, 2.6 mM sodium pyrophosphate, 20.4 mM Tris-HCl and 1% (v/v) Triton X-100, supplemented with 0.1 mM phenylmethanesulfonylfluoride (PMSF), 12% (v/v) protease inhibitor cocktail (Sigma #P8340), 1 mM dithiothreitol (DTT) and 20 mM sodium fluoride. The protein content of each sample was determined by the Bradford method[54]. Laemmli buffer 6x (composed by 188 mM Tris-HCl, 2.2 mM sodium dodecyl sulfate (SDS), 0.03% (w/v) bromophenol blue and 79% (v/v) glycerol) was added 50 μg of protein samples that were then denatured at 95°C for 5 min. Samples containing 50 μg protein were separated by 12% polyacrylamide gel electrophoresis and electrophoretically transferred to a polyvinylidene difluoride (PVDF) membrane. Membranes were then stained with Ponceau S (P3504, Sigma Aldrich), a reliable method for total protein normalization in western blotting[55]. After membrane blocking with 5% nonfat dry milk (Bio-Rad) in Tris-Buffered Saline Tween (TBS-T; 50 mM Tris-HCl, pH 8; 154 mM NaCl and 0.1% (v/v) Tween 20) for 2 h at room temperature under continuous stirring, membranes were incubated overnight at 4°C, under stirring, with the primary antibodies against GPx-1/2 (1:1000, sc-133152, RRID: AB_2112240, Santa Cruz), catalase (1:1000, sc-271803, RRID: AB_10708550, Santa Cruz), glutathione reductase (1:1000, sc-133136, RRID: AB_2115652, Santa Cruz), SOD-2 (1:1000, sc-133134, RRID: AB_2191814, Santa Cruz), SOD1 (1:2000, sc-11407, RRID: AB_2193779, Santa Cruz) or actin (1:5000, MAB1501, RRID: AB_2223041, Millipore). Once incubation was complete, membranes were incubated with horseradish peroxidase (HRP)-conjugated secondary antibodies goat anti-rabbit (1:1000, #7074, RRID: AB_2099233, Cell Signaling) and mouse anti-goat (1:750, sc-2354, RRID: AB_628490, Santa Cruz) for 1 h at room temperature under continuous agitation. Membranes were incubated with Enhanced Chemiluminescence reagent (ECL, #1705061, Bio-Rad) and imaged using a VWR^®^ Gel Documentation System Imager Chemi 5QE (VWR, USA). Band density was measured using the TotalLab TL120v2008 software (Newcastle, UK) and the results were expressed as % of control of the same cohort.

### 2.6 Evaluation of intracellular oxidative stress levels by H_2_DCFDA oxidation assay

Oxidative stress levels in lymphoblasts were evaluated through indirect measuring of 2’,7’-dichlorodihydrofluorescein diacetate (H_2_DCFDA) oxidation by flow cytometry, based on the assumption that DCF fluorescence is proportional to the concentration of reactive redox species present in the cytoplasm[56]. The volume of cell suspension containing 2 x 10^6^ cells was centrifuged at 259 g, at room temperature, for 5 min. The supernatant was aspirated and resuspended with 600 μL of phosphate-buffered saline (PBS, containing 137 mM NaCl, 2.7 mM KCl, 1.4 mM K_2_HPO_4_, and 4.3 mM KH_2_PO_4_, at pH 7.4) and divided in two microtubes, each with 1 x10^6^ cells/ 300 μL PBS. Afterwards, cell suspensions were incubated in the dark at 37°C in the absence or presence of 5 μM H_2_DCF-DA dye, for 15 min. After incubation, the samples were placed on ice to stop H_2_DCF-DA loading. Approximately 50,000 gated fluorescence events were acquired using an Accuri^™^ C6 flow cytometer (Becton Dickinson) with an excitation filter of 530/30 nm, and analysis was performed with FlowJo V10 software (Ashland, OR, USA). DCF fluorescence was expressed as % of the pool of controls.

### 2.7 Evaluation of lipid peroxidation

Lipid peroxidation was assessed by the fluorimetric determination (excitation at 515 nm and emission at 553 nm; FP-2020/2025, Jasco, Tokyo, Japan) of malondialdehyde (MDA) adducts separated by high-performance liquid chromatography (HPLC; Gilson, Lewis Center, Ohio, USA) using the ClinRep complete kit (RECIPE, Munich, Germany). The results are expressed as μmol of MDA per mg of protein (μmol/mg prot).

### 2.8 Evaluation of t-BHP cytotoxicity

Resazurin reduction assay was used to assess cell viability. To determine the cytotoxic effects of t-BHP, 160,000 cells/well were seeded in 96-well plates, with a final volume of 50 μL per well and treated with t-BHP (0, 250, 500, 5000 μM). After 6 h, 50 μL of resazurin solution at 20 μg/mL was added and incubated for 2 h at 37 °C and 5% CO_2_. The reduction of resazurin to resorufin was measured fluorometrically at 540 nm excitation and 590 nm emission in a Cytation 3 microplate reader (BioTek Instruments Inc., USA). The results were expressed as % of untreated cells, for each cell line.

### 2.9 Glutathione peroxidase (GPx) activity

GPx is an antioxidant enzyme responsible for eliminating hydrogen peroxide (H_2_O_2_), using reduced glutathione (GSH)[57]. GPx activity was determined using a colorimetric method [58]. Lymphoblasts were seeded at 400,000 cells in T75 flask containing 60 mL RPMI medium and incubated for 48 h to reach a cell density of around 1×10^6^ cells/mL in each assay. Then, cells were harvested, lysed and protein was quantified using the BCA method[53]. Cell lysates containing 30 μg protein were incubated with 15 μL of transforming solution (composed of 4.5 mM KCN, 0.45 mM K_3_[Fe(CN)_6_] and 0.25 mM KH_2_PO_4_) and incubated for 5 min. Then, each well of a 96-well plate was loaded with 40 μL H_2_O milliQ, 10 μL sample (33 μg protein), 10 μL 10 mM GSH, 10 μL GR 2.4 U/mL, 10 μL phosphate buffer 0.25 M (K_2_HPO_4_, KH_2_PO_4_), 10 μL 12 mM tert-butyl hydroperoxide (t-BHP) and a baseline reading at 340 nm, 37°C for 2 min was performed. Finally, 10 μL 2.5 mM NADPH was added, and the oxidation of NADPH was measured at 340 nm, 37°C, for 5 min in a VICTOR 3 microplate reader (Perkin Elmer). The results were expressed as % of the pool of controls.

### 2.10 Glutathione reductase (GR) activity

GR is an antioxidant enzyme that reduces glutathione, thus maintaining the pool of this cellular antioxidant[57]. GR activity was determined using a colorimetric method based on the oxidation of NADPH[58]. Lymphoblasts were seeded at 400,000 cells in T75 flasks containing 60 mL RPMI medium and incubated for 48 h to reach a cell density of around 1×10^6^ cell/mL in each assay. Then, cells were harvested, lysed and protein was quantified using the BCA method[53].

Each well of a 96-well plate was loaded with 90 μL of 0.12 M phosphate buffer pH 7.2, 5 μL sample (25 μL protein), 5 μL of 15 mM (EDTA) and 5 μL of 65.3 mM oxidized glutathione (GSSG). Then, the plates were incubated for 3 min at 37°C, followed by baseline reading at 340 nm, at 37°C for 3 min. Finally, 5 μL 4.8 mM NADPH was added, and the oxidation of NADPH was measured at 340 nm, at 37°C for 5 min in a VICTOR 3 microplate reader (Perkin Elmer). Results were expressed as % of the pool of controls.

### 2.11 Intracellular glutathione levels

Total glutathione content was determined by the glutathione reductase enzymatic method [18]. Briefly, lymphoblasts were lysed with assay buffer containing 0.1% (v/v) Triton X-100 and 0.6% 5-sulfosalicylic acid and disrupted by sonication. Cell extracts were centrifuged at 3,000 g at 4°C for 5 min. Supernatants were then used for quantification of oxidized glutathione (GSSG) and total glutathione (GSH + GSSG) levels. For GSSG determination, samples were pre-incubated with 2-vinyl-pyridine for 1 h at room temperature, followed by triethanolamine for 10 min. Glutathione standards were run simultaneously with the samples. Reaction was initiated by adding 120 μL of working solution (5,5’-dithiobis 2-nitrobenzoic acid and GR with 20 μL of cell extract (50 μg protein) or glutathione standards. Next, 60 μL of 0.8 mM NADPH solution was added and the increase in absorbance was recorded at 412 nm wavelength for 2 min. GSH and GSSG values obtained from the standard curves were expressed as nmol/mg protein.

### 2.12 Catalase activity assay

Catalase activity was assessed using a Catalase Activity Kit (#EIACATC; Invitrogen, Hampton, USA), according to the manufacturer’s instructions. Briefly, pellets containing 30 x 10^6^ cells were rinsed with the provided Assay Buffer, sonicated, and centrifuged at 300 x g for 15 min at 4°C. Then, 25 μL supernatant, corresponding to 0.5 μg protein, and catalase standards, were added to H_2_O_2_ reagent in 96-well plates and incubated for 30 min, followed by substrate and HRP solution and incubation for 15 min at room temperature. Increased levels of catalase activity in the samples decrease H_2_O_2_ concentration observed during the reduction of the colorimetric detection reagent into a pink-colored product. Absorbance was then read at 560 nm using a BioTek^®^ Cytation3^™^ microplate reader (BioTek, Agilent, Winooski, VT, USA). Catalase activity was calculated for all samples according to a standard curve of known catalase activities. Results were expressed as % of the pool of controls.

### 2.13 Measurement of vitamin E levels

Cell lysates were prepared as described for GPx activity assessment. Then, 500 μL cell lysate with 1 mg/mL protein were added to 1.5 mM of 10 mM SDS, 2 mL of ethanol and 2 mL of n-hexane, and lipids were extracted from the upper phase formed after centrifugation. Vitamin E (vit E) levels were quantified by reverse-phase HPLC using an analytic column Spherisorb S10w (250 mm × 4.6 mm), eluted at 1.5 mL/min with n-hexane modified with 0.9% methanol, and detected in a spectrophotometer (Gilson, Lewis Center, Ohio, USA) at 287 nm[59]. The sample volume injected (20 μL) corresponded to 2 μg protein. A calibration curve made with known concentrations of α-tocopherol was used to calculate the concentration of vitamin E present in the extracts, and results were expressed as μM.

### 2.14 Computational data analysis

Orange 3.27.1 [60] was used for the computational data analysis and visualization. Features were ranked according to the chi-square scores between each individual feature and the target ‘Experimental Group’ (Control, undSOD1, mutSOD1). The top 7 features were further considered for their potential to distinguish samples according to the target. A linear projection was generated for the selected features after applying principal component analysis (PCA) for dimensionality reduction. The separation between the experimental classes, provided by the selected features, was also visualized using a heatmap coupled with hierarchical clustering analysis, using Seaborn 0.11.1[61].

### 2.15 Statistics

Paired and pooled statistical analyses were performed using GraphPad Prism 8.02 software (GraphPad Software, Inc., San Diego, California, USA). Data are represented in bar graphs (mean±SEM) with dot plots for the number of experiments indicated in figure legends, or in dots (mean±SEM) with adjusted mathematical curves. Statistical significance was set at P<0.05 and determined by the Kruskal Wallis method, followed by Dunn’s multiple comparisons test or by two-way ANOVA, as appropriate.

## 3 Results

We first analyzed their proliferation to characterize the selected lymphoblast cell lines and set up the conditions for the following experiments. Cells were seeded at similar concentrations, in suspension, and counted after 24, 48, 72 or 96 h. We compared proliferation in cells from the same cohort (Fig.1 a-c) and in cells belonging to the same experimental group (Fig.1 d-f). For C1 (Fig.1 a), undSOD1 cells presented significantly increased proliferation at 72 and 96 h, compared to the control group, while for C2 (Fig.1 b), both undSOD1 and mutSOD1 lymphoblasts presented higher proliferation than the controls, for the same time points. The opposite effect was found in C3 (Fig.1 c), with undSOD1 and mutSOD1 presenting lower proliferation compared to control cells, while undSOD1 lymphoblasts showed significantly lower cell numbers than mutSOD1 at 72 h. Differences in proliferation unrelated to the experimental groups, possibly due to sex, age-of-onset and/or type of SOD1 mutation, became evident when we compared cells belonging to the same experimental group (Fig.1 d-f). Among the different controls, lymphoblasts from the youngest cohort (C3) presented the highest proliferation, significantly higher than C1 after 72 h, and significantly higher than C2 after 48 h (Fig.1 d). Control lymphoblasts from C2 showed the lowest proliferation, being significantly lower than C1 after 72 h (Fig.1 d). In the undSOD1 group, lymphoblasts from C2 and C3 presented lower proliferation than C1 after 72 h (Fig.1 e). The profiles were more similar between cohorts in mutSOD1 cells, with C3 having significantly more cells at 72 and 96 h than C2, while C2 showed significantly less cells at 96 h than C1 (Fig.1 f).

**Figure 1:**
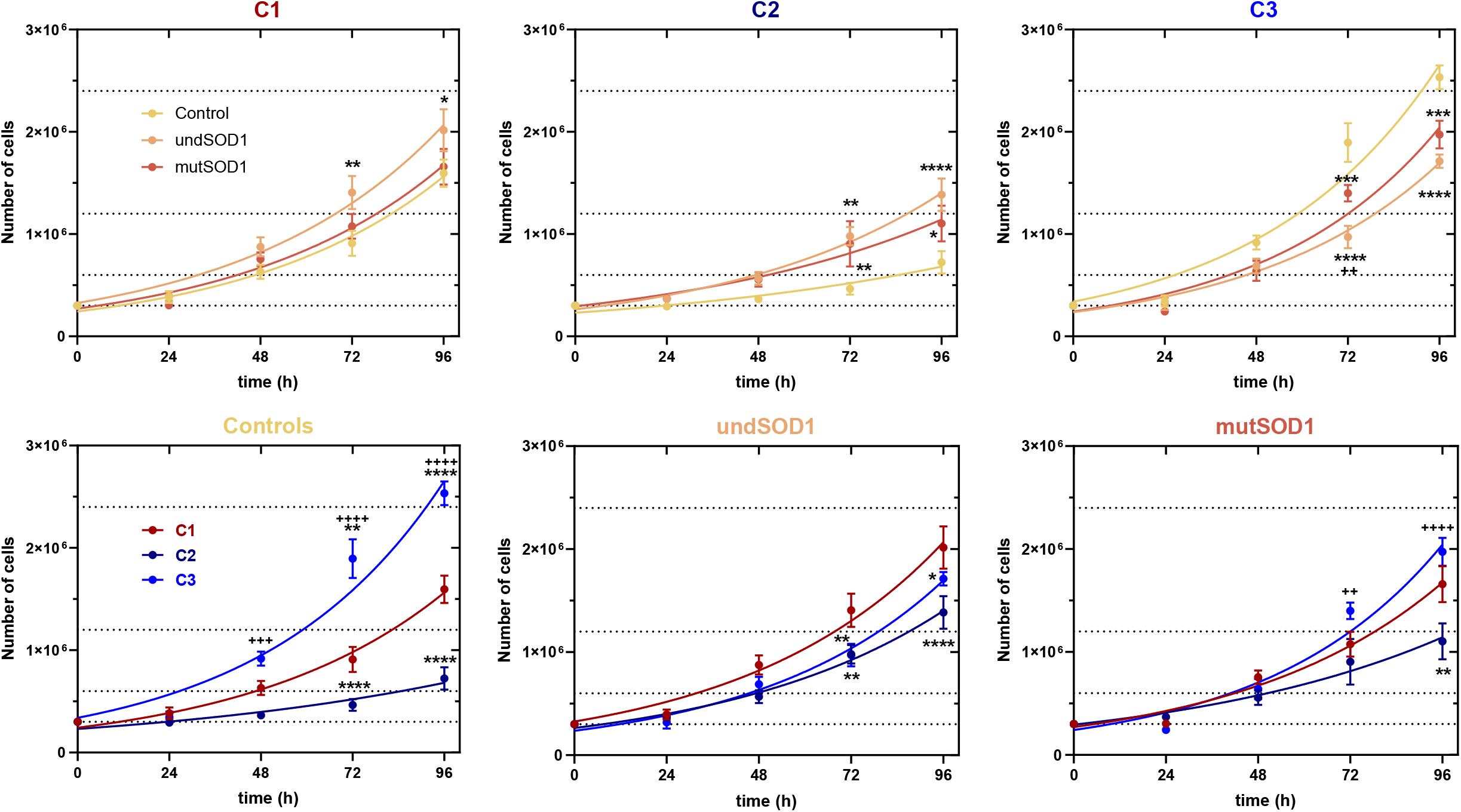
Comparison between growth curves obtained for lymphoblasts from different (a-c) cohorts or (d-f) experimental groups, plated at 30,000 cells/condition (at t0). a) C1-Females 46 years old, b) C2-Males 46 years old, c) C3-Males 25/26 years old, d) Healthy controls, e) undSOD1, f) mutSOD1. Exponential curves were adjusted to the data, and doubling times were calculated as 35.7 h for Control_C1, 36.0 h for undSOD1_C1, 36.5 h for mutSOD1_C1, 61.4 h for Control_C2, 39.8 h for undSOD1_C2, 49.0 h for mutSOD1_C2, 32.3 h for Control_C3, 33.8 h for undSOD1_C3, and 31.2 h for mutSOD1_C3. The data was compared by Two Way ANOVA and Tukey post-test, results expressed as Mean ± SEM, n≥4; ****P < 0.0001, ***P < 0.001, **P < 0.01, *P < 0.05 vs. C1; ++++P < 0.0001, ++P < 0.01 vs. C2. The datasets related to this figure are available at https://doi.org/10.6084/m9.figshare.19146161

Next, we assessed SOD activity, gene, and protein expressions in lymphoblasts obtained from ALS patients with mutSOD1or undSOD1 and respective age- and sex-matched controls (Fig. 2). mutSOD1 cells from C1 presented a significant decrease in SOD activity (Fig.2 a) and in SOD1 protein levels (Fig.2 d). However, no significant changes in SOD1 or SOD 2 transcripts (Fig.2 b-c) or in SOD2 protein levels (Fig.2 e) were observed. mutSOD1 cells from C3 also presented a significant decrease in SOD activity (Fig.2 a).

**Figure 2:**
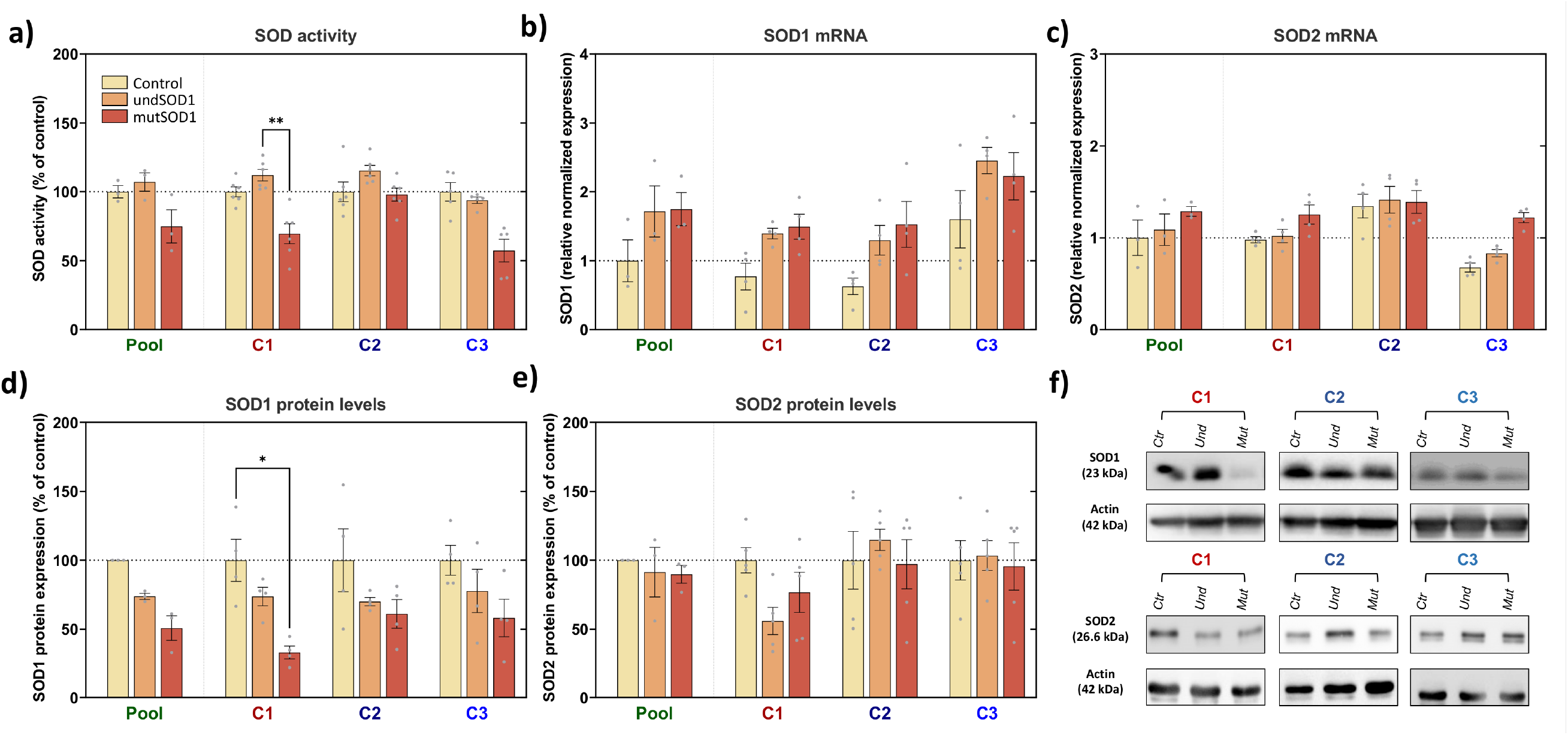
Superoxide dismutase (SOD) activity and expression in lymphoblasts obtained from ALS patients with mutant SOD1 (mutSOD1) and unknown SOD1 mutations (undSOD1). Cells were plated and treated as described in Materials and Methods. a) Total SOD activity was analyzed using a commercial kit and normalized in percent of the respective control. b-c) Gene expression was evaluated by qPCR, corrected to the geometric mean of TBP, YWHAZ, PUM1 and B2M and normalized to the pool of control samples. d-e) protein expression was corrected to actin in the same sample and normalized in percentage of the respective control. Data was compared by the Kruskal Wallis method, followed by Dunn’s multiple comparisons test. Results are the mean±SEM from at least 3 independent experiments. *p<0.05, **p<0.01, ***p<0.01 compared to respective control. +p<0.05, ++p<0.01 compared to respective undSOD1 lymphoblasts. The datasets related to this figure are available at https://doi.org/10.6084/m9.figshare.19146332

We next assessed markers of oxidative stress-induced damage and response. We measured H_2_DCFDA oxidation as an indicator of intracellular levels of reactive redox species, and thus a marker of oxidative stress (Fig.3 a). This marker was increased in mutSOD1 from C1, compared to the respective control (Fig.3 a), which is negatively correlated with the changes observed in SOD1 protein levels and total SOD activity in this cohort (Fig.2 c-d). In addition, oxidative stress also increased in undSOD1 and mutSOD1 lymphoblasts from C3 compared to the control. No changes in lipid peroxidation were observed in any experimental groups or cohorts, as analyzed MDA levels (Fig.3 b). Besides evaluating cellular oxidative stress markers under basal conditions, we next analyzed the lymphoblast metabolic activity after incubation with t-BHP (Fig.3 d-f). In C1, significant increases in oxidative stress susceptibility were observed in undSOD1 and mutSOD1 lymphoblasts, compared to controls (Fig.3 d). In C2, the only significant result was the increased susceptibility of undSOD1 lymphoblasts to intermediate concentrations of t-BHP, compared to mutSOD1 lymphoblasts (Fig.3 d). In C3, only mutSOD1 cells presented increased susceptibility to the higher concentration of t-BHP compared to controls (Fig.3 f).

**Figure 3:**
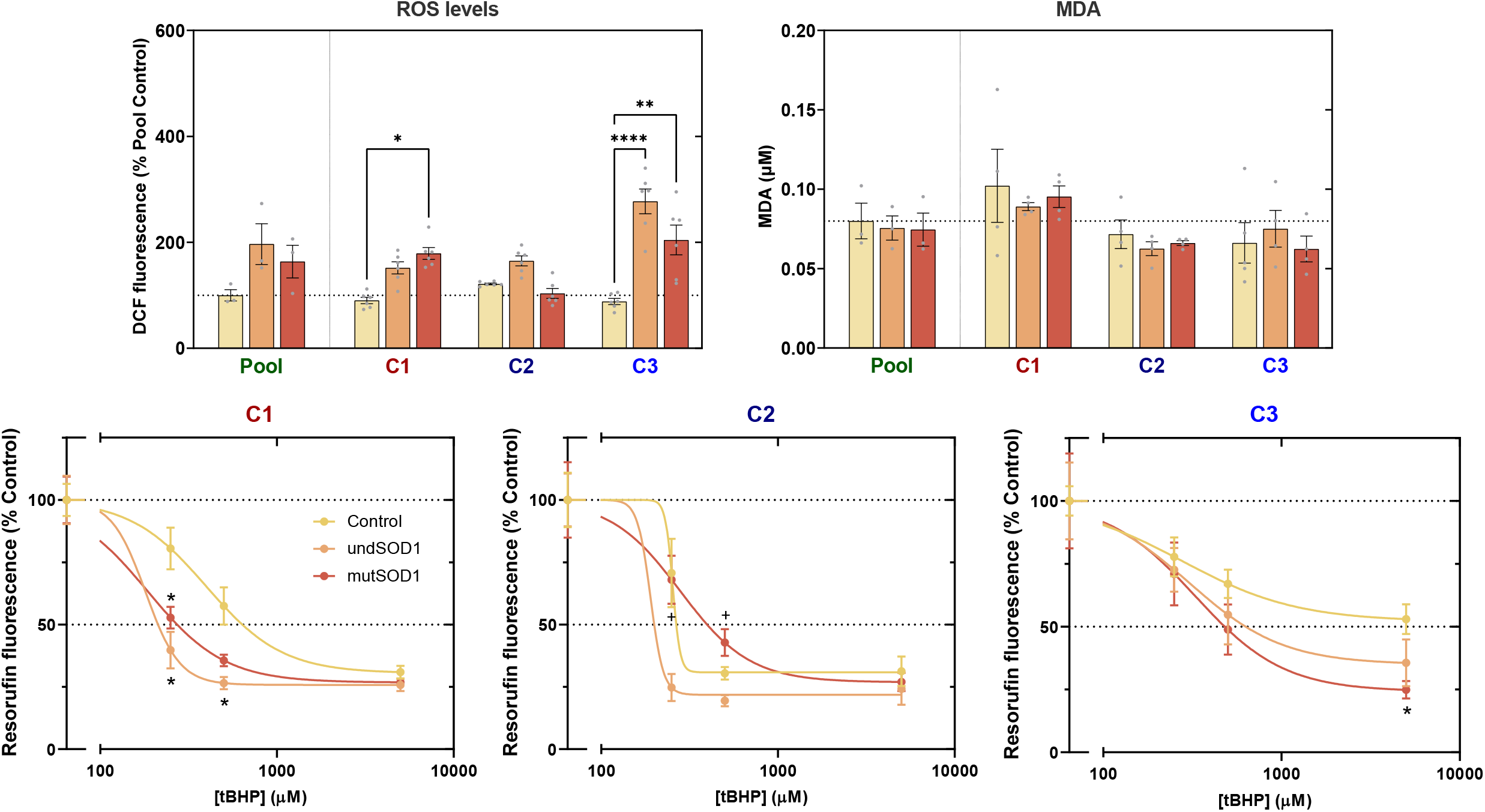
Oxidative stress markers and metabolic response to oxidative stress (t-BHP) in lymphoblasts obtained from ALS patients with mutant SOD1 (mutSOD1) and unknown SOD1 mutations (undSOD1). Cells were plated and treated as described in Materials and Methods. a) Intracellular oxidative stress was evaluated by flow cytometry using the H_2_DCFDA assay. b) Malondialdehyde (MDA) levels were assessed by HPLC as a marker of lipid peroxidation. Data was compared by the Kruskal Wallis method, followed by Dunn’s multiple comparisons test. c-e) Cellular susceptibility to oxidative damage by increasing concentrations of tert-butyl-hydroperoxide (t-BHP) was analyzed in each cohort by following metabolic activity using the resazurin reduction assay, and four-parameter logistic curves were adjusted to the data, which was compared by Two Way ANOVA and Tukey post-test. Results are the mean±SEM from at least 3 independent experiments. *p<0.05, **p<0.01, ****p<0.001 compared to respective control. ++p<0.01 compared to respective undSOD1 lymphoblasts. The datasets related to this figure are available at https://doi.org/10.6084/m9.figshare.19146386

We then assessed antioxidant defenses associated with glutathione redox cycling. No significant changes were observed in GPx1 or GPx4 transcripts (Fig.4 a,b), or in GPx protein levels (Fig.4 c,d) or activity (Fig.4 e). For GR activity (Fig.4 f), undSOD1 from C1 showed higher GR activity compared to the respective control, and in C2, undSOD1 had higher GR activity than mutSOD1 (Fig.4 f), despite no changes were found in GR protein levels (Fig.4 g-h). No statistically significant changes were observed in GSH or GSSG levels, but the ratio GSH/GSSG showed a tendency to be decreased in undSOD1 and mutSOD1 from C1 and C3, compared to the respective controls. The alterations in this ratio seem to have different origins, as there was a non-significant decrease in GSH levels in undSOD1 (Fig. 4 i), whereas there was a non-significant increase in GSSG levels in mutSOD1 (Fig. 4 j).

**Figure 4:**
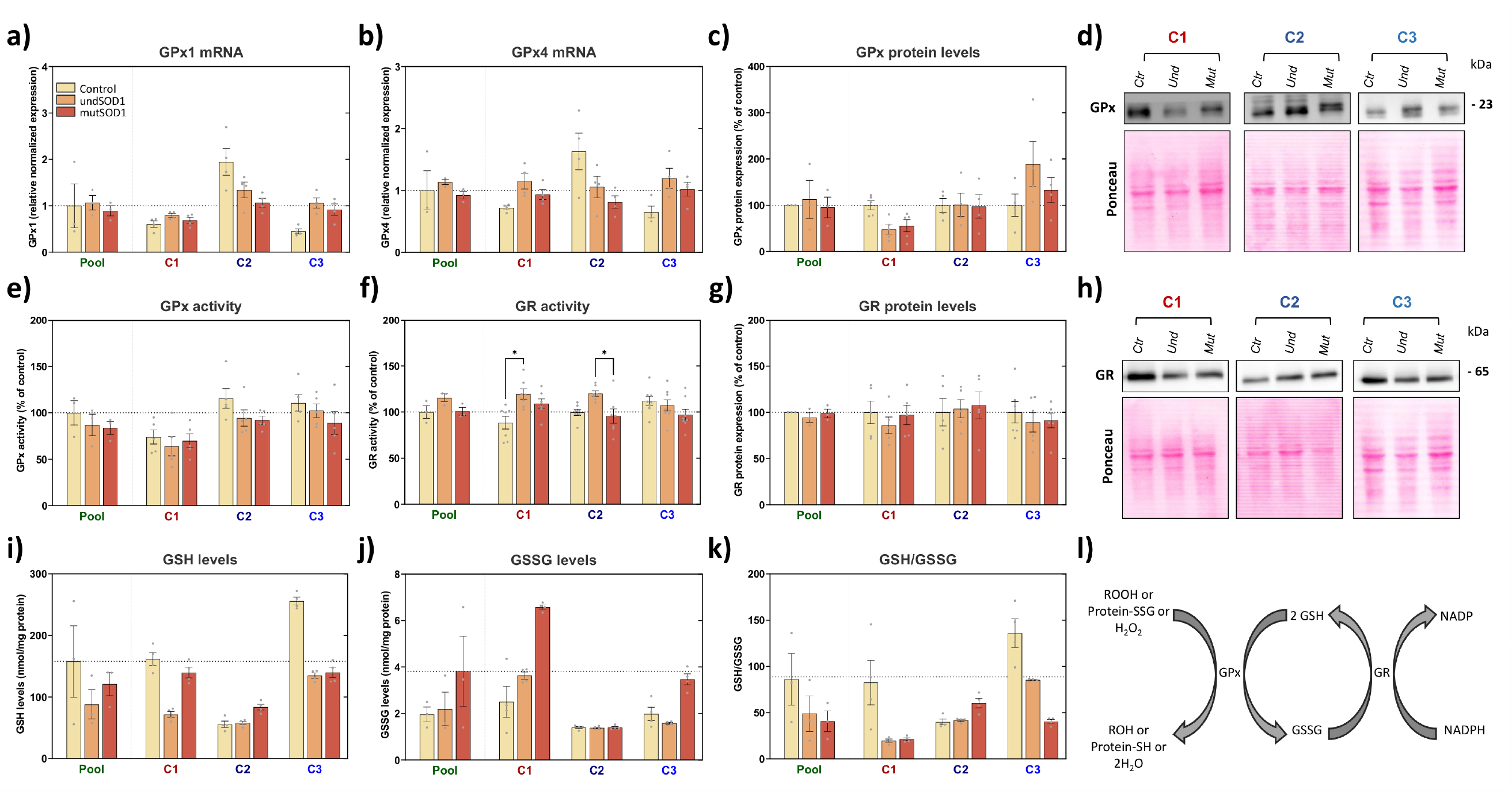
Analysis of glutathione redox cycle markers in lymphoblasts obtained from ALS patients with mutant SOD1 (mutSOD1) and unknown SOD1 mutations (undSOD1). Cells were plated and treated as described in Materials and Methods. a-b) Glutathione peroxidase (GPx) transcript levels were analyzed by qPCR. e) GPx and f) Glutathione reductase (GR) activities were analyzed using spectrophotometric assays. c-d) GPx and g-h) GR protein levels were analyzed by western blotting. i) GSH, j) GSSG and k) GSH/GSSG levels were evaluated by HPLC. l) glutathione redox cycle. Data was compared by the Kruskal Wallis method, followed by Dunn’s multiple comparisons test. Results are the mean±SEM from at least 3 independent experiments. *p<0.05, **p<0.01, ***p<0.01 compared to respective control. +p<0.05, ++p<0.01 compared to respective undSOD1 lymphoblasts. The datasets related to this figure are available at https://doi.org/10.6084/m9.figshare.19146437

We further assessed other intracellular antioxidant defenses, including catalase, vitamin E and the master antioxidant response transcription factor Nrf2 (Fig.5). Catalase mRNA levels were only significantly increased in undSOD1 and mutSOD1 from C2, compared to the respective control (Fig.5 a), accompanied by a non-significant increase in catalase protein levels (Fig.5 b-c). However, catalase activity was only significantly increased in undSOD1 from C2, compared to the respective mutSOD1 (Fig.5 d). These changes may be correlated with the other altered parameters found in C2 lymphoblasts from ALS patients (undSOD1 and mutSOD1), namely the increased proliferation rate (Fig. 1b) and the differential susceptibility to t-BHP (250-500 μM) (Fig. 3d). No significant changes were observed in vitamin E (Fig.5 e) or in Nrf2 mRNA (Fig.5 f) levels.

**Figure 5:**
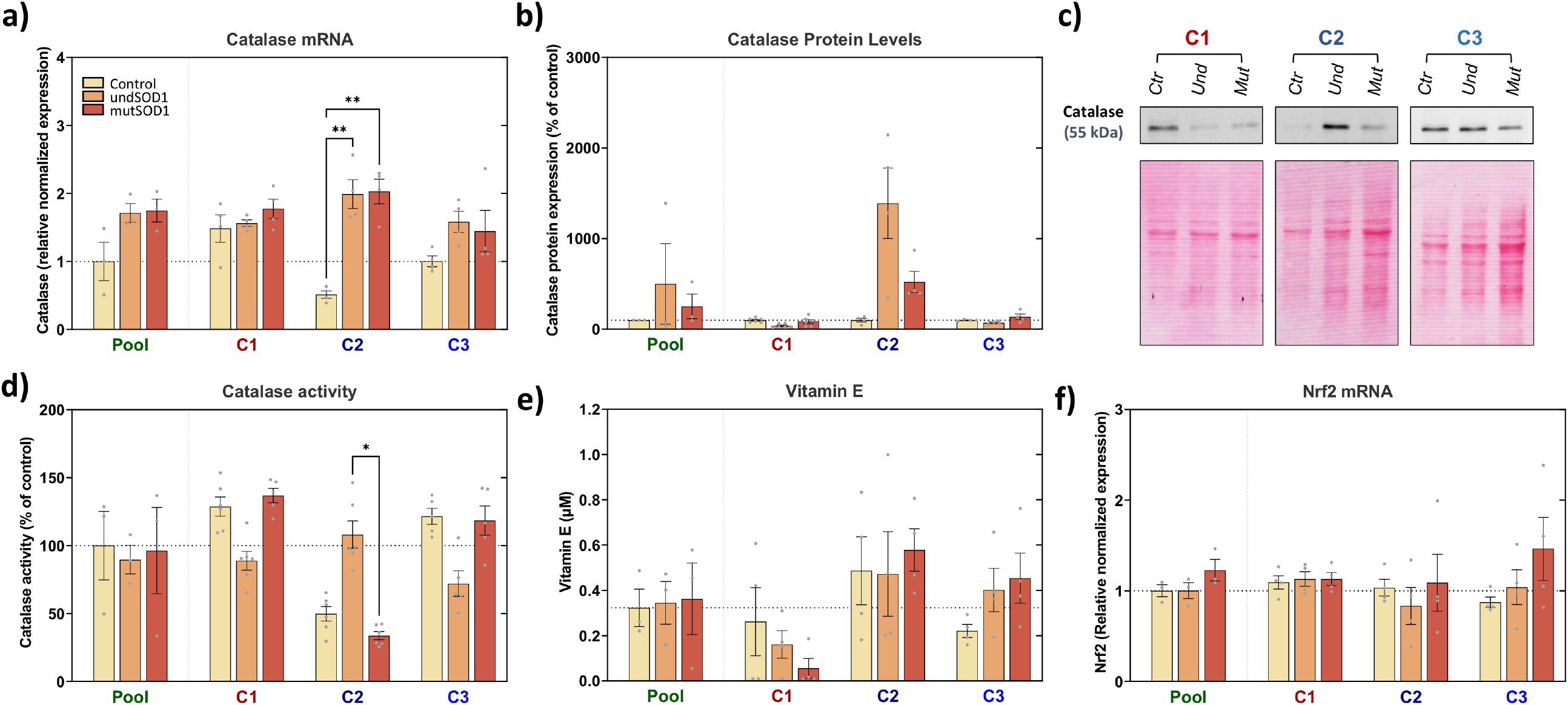
Analysis of key antioxidant defenses in lymphoblasts obtained from ALS patients with mutant SOD1 (mutSOD1) and unknown SOD1 mutations (undSOD1). Cells were plated and treated as described in Materials and Methods. a,f) Catalase and Nrf2 transcript levels were analyzed by qPCR. b-c) Catalase protein levels were analyzed by western blotting. e) Catalase activity was analyzed using a commercial kit. e) Vitamin E levels were analyzed by HPLC. Data was compared by the Kruskal Wallis method, followed by Dunn’s multiple comparisons test. Results are the mean±SEM from at least 3 independent experiments. **p<0.01, ***p<0.01 compared to respective control. The datasets related to this figure are available at https://doi.org/10.6084/m9.figshare.19146482

Since no significant differences were found in the pooled analysis, other strategies were needed to try and find hidden patterns in the data. Thus, we performed multidimensional data analysis to integrate all the results and identify which alterations are more characteristic of undSOD1 or mutSOD1 lymphoblasts (Fig.6). All the dataset features were ranked according to their chi-square scores to identify the most discriminant features (Fig.6 a). For the features previously normalized to the respective control (those related with western blotting), the raw (non-normalized) values were used, to avoid introducing an artificial similarity between controls belonging to different cohorts. This analysis evidenced 7 features with high discriminative power regarding the target variable (experimental group: control, undSOD1, mutSOD1), including total SOD activity, intracellular oxidative stress (assessed by DCF fluorescence), catalase mRNA levels, SOD1 protein levels (raw values), Nrf2 mRNA levels, resazurin reduction in the presence of 5000 μM t-BHP (% of untreated cells) and GR activity. These features were used for subsequent clustering (Fig.6 b) and PCA (Fig.6 c) analyses. The analyses revealed a perfect separation of samples from control and ALS (undSOD1, mutSOD1) samples (Fig.6 b-c). The ALS samples were also well separated into two clusters, corresponding to undSOD1 and mutSOD1, both in the cluster map (Fig.6 b), and in the PCA (Fig.6 c).

**Figure 6:**
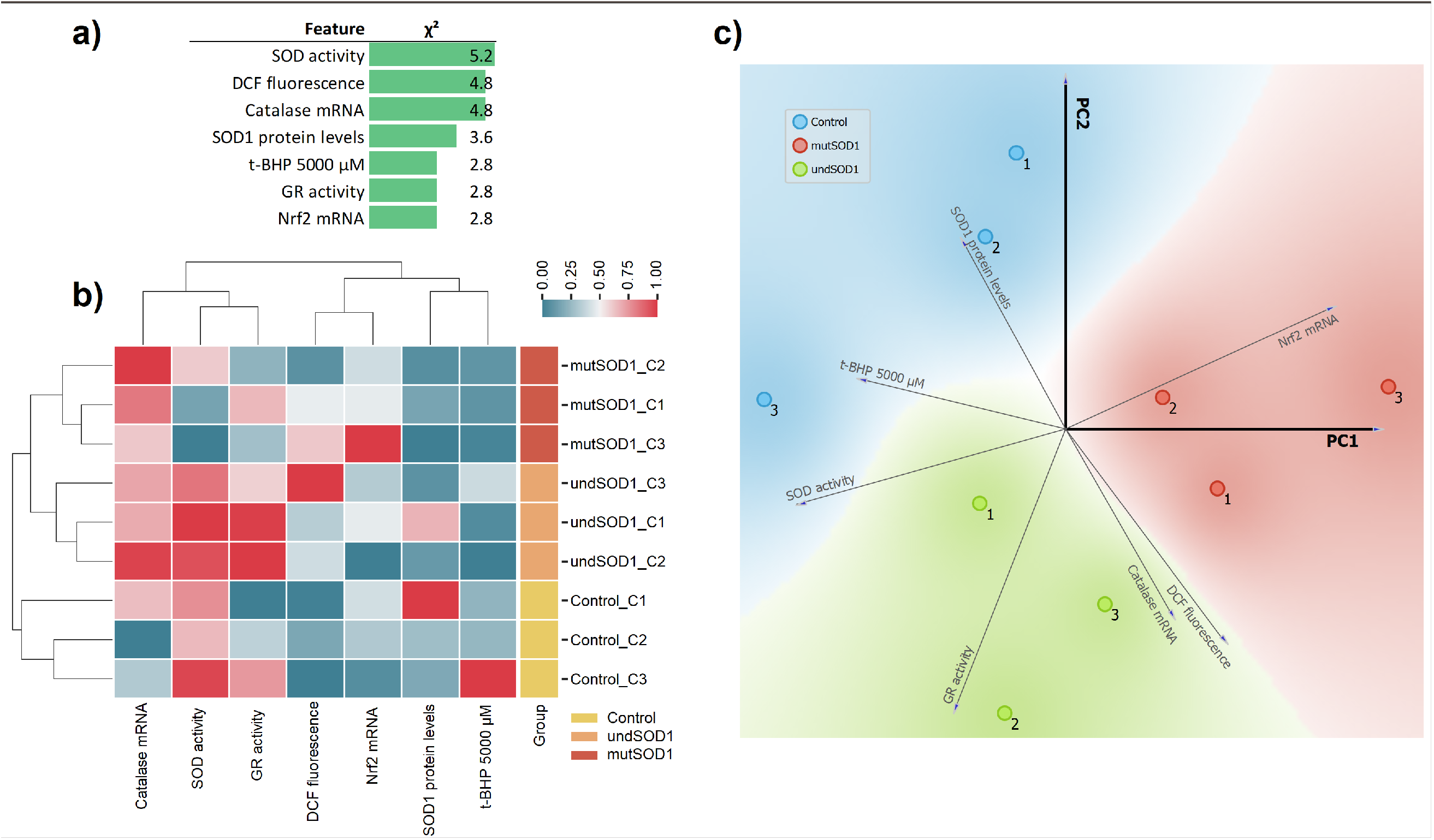
Integrative analysis of oxidative stress profiles of lymphoblasts obtained from ALS patients with mutant SOD1 (mutSOD1) and unknown SOD1 mutations (undSOD1). a) Features were ranked according to their chi-square scores, evidencing the features with more discriminative power between experimental conditions and enabled the selection of the 7 most discriminant features for subsequent b) clustering and c) principal component (PCA) analyses. The datasets related to this figure are available at https://doi.org/10.6084/m9.figshare.19146491

## 4 Discussion

The present work characterized oxidative stress fingerprints in lymphoblasts from ALS patients with (mutSOD1) or without (undSOD1) known SOD1 mutations, compared with age- and sex-matched controls, comprising 46-year-old females (C1), 46-year-old males (C2) and 26/27-year-old males (C3). Our results were highly heterogeneous between cohorts of different age and sex, which is not surprising due to the intrinsic heterogeneity of ALS, which presents different rates of progression, patterns of spread, various sites of onset between patients[62], association to various genetic mutations[63] and, in most cases, with no identified mutations. Thus, some degree of heterogeneity among the redox state of circulating blood cells is also expected between patients. We also cannot exclude some heterogeneity introduced by the lymphoblast transformation process. Due to the intrinsic disease heterogeneity, no single treatment strategy is expected to fit all patients, being essential to identify small responder groups within larger patient populations[64, 65], which can be potentially achieved by studying the oxidative stress profile of each patient, helping to develop personalized therapeutic strategies. This can potentially be done using peripheral blood mononuclear cells (PBMCs), which enable measuring multiple biochemical parameters, even from cryopreserved cells[32, 66]. However, the amount of sample obtained from a given patient is always extremely limited, which is even more critical for diseases like ALS, a rare and rapidly progressing fatal orphan disease (ORPHA:803), limiting the number of patient samples obtained for analysis. An alternative is the use of lymphoblasts, which are immortalized PBMCs that can be obtained from biobanks such as the Coriell Institute. These cells proliferate in culture, enabling measuring multiple functional cellular parameters at different time points, and have also been proved helpful in studying mitochondrial function in ALS [67] and other neurodegenerative diseases[68–70], as they can be manipulated and challenged *in vitro* to highlight subclinical alterations.

mutSOD1 lymphoblasts were previously used to demonstrate the existence of hyper oxidized SOD1 with toxic properties in ALS patient-derived cells and allowed the identification of common SOD1-dependent toxicity between mutant SOD1-linked familial ALS and a subset of sALS[71]. This highlights how lymphoblasts can provide opportunities for biomarker development, sub-classifying ALS, and designing more disease-modifying therapies[71]. Moreover, increased DCF fluorescence was reported in lymphoblasts from familial ALS patients with SOD1 mutations compared with sporadic ALS and normal controls (spouses of ALS patients), although the DCF oxidation levels were not directly correlated with SOD1 activity[72]. We investigated several oxidative stress hallmarks in lymphoblasts from each ALS patient in the present work. Although a high heterogeneity was found, we identified specific alterations that may have a diagnostic and therapeutic interest.

Importantly, most data correlating oxidative stress markers with ALS progression has been obtained using mutSOD1 animal models, in which the mechanisms behind oxidative stress are easy to devise but which are a surrogate for only a small proportion of all ALS patients. Non-SOD1 related ALS, which represents the highest number of patients among other familial ALS mutations and non-SOD1-related sALS, has been less studied, due to the lack of proper animal models, although environmental risk factors suggest that oxidative stress may be related to sALS onset. In sALS patients, deregulation of the cellular redox systems has been suggested by higher free radical levels found in the blood and serum of patients[32, 73], and evidence of oxidative damage includes the increase in protein carbonyls[34, 35, 74, 75], 8-OHdG[47, 49], MDA-modified proteins[34], 4-HNE protein conjugates[36, 76], and nitrotyrosine products[77–79] in the spinal cord. Moreover, erythrocytes from sALS patients presented an increase in lipid peroxidation associated with a decrease in catalase, GR, and glucose-6-phosphate dehydrogenase activities and a decrease in GSH levels, especially in cases with longer disease duration times were measured[80], suggesting a correlation between these parameters and the duration of ALS. Similarly, lower activities of GPx and GR were found in erythrocytes from sALS patients with a faster disease progression rate, and the plasma total antioxidant status was also decreased in the same group of patients[81]. Thus, there seems to be a correlation between oxidative stress and sALS.

In ALS patients, more than 180 mutations have been identified across the coding and non-coding regions of the SOD1 gene [52, 82]. The influence of these mutations on dismutase activity is considerably variable, having been associated with decreased[38], maintained[83, 84], or increased[38, 85] enzymatic activity, compared to wild-type SOD1. Thus, it has been proposed that mutSOD1 exerts its deleterious effect by a toxic gain of function rather than by altered SOD1 activity, given the lack of correlation between SOD1 dismutase activity and aggressiveness of clinical phenotypes[86] and since SOD1 knockout mice do not develop the ALS phenotype per se[87]. mutSOD1 has been associated with a loss of redox homeostasis and excessive production of reactive redox species[23, 72, 88–91], and increased levels of reactive redox species (mainly, H_2_O_2_) were previously found in lymphoblasts of fALS cases with mutSOD1[72]. Nonetheless, mutations in SOD1 may destabilize the protein structure, resulting in misfolded SOD1, which in its soluble form, binds to intracellular organelles (e.g. mitochondria) disrupting their function, or forms toxic insoluble aggregates[1]. Indeed, association of misfolded SOD1 with mitochondria in spinal cord of both SOD1G93A rats and SOD1G37R mice was found to occur prior to disease onset and was correlated with increased superoxide production and mitochondrial damage[23, 88].

Mitochondrial mutSOD1 was previously associated with respiratory chain impairment and to a shift in the mitochondrial redox balance, leading to decreased GSH/GSSG ratio[89]. Decreased levels of GSH and increased levels of GSSG were reported in NSC34 motor neuron-like cells and lumbar tissues of mutant SODG93A mice spinal cord[90]. Indeed, the decrease in total glutathione levels (GSH and GSSG) in spinal cord mitochondria from hSOD1G93A mice was linked to decreased survival, possibly due to compromised mitochondrial function and morphology, increased oxidative stress, astrogliosis and motor neuron loss[91].

Previous works reported normal catalase activity in erythrocytes from sporadic and familial ALS[92]. However, a later study found a significant decrease in catalase activity in erythrocytes of sALS patients compared to controls, whereas catalase activity in plasma and serum was similar in both groups[93]. Moreover, blood and CSF vitamin E levels were unaltered in ALS[94, 95].

Toxic gain of function of mutSOD1 was suggested to perturb the activation of the nuclear factor erythroid-2-related factor 2 (Nrf2) pathway[96], a major regulator of the phase II antioxidant response, including GPx, catalase, GR[6] and even SOD1[97]. Indeed, Nrf2 expression decreased in NSC-34 motor neuron-like cells expressing mutSOD1 and motor neurons isolated from familial SOD-associated ALS patients[32, 98]. In line with this, activation of the Nrf2-antioxidant response element (ARE)[99] oxidative stress responsive system occurs in distal muscles prior to the disease onset in mutSOD1 mice[100], supporting the hypothesis that increased muscle oxidative stress is involved in an early phases of ALS, eventually followed by axonal “dying back” and culminating with loss of motor neurons. In our work, no alterations were found in Nrf2 transcript levels (Fig. 4 f), although we cannot exclude changes in its protein levels and/or activity.

Following conventional data analysis, no common changes between mutSOD1 vs control and undSOD1 vs control in the different cohorts were evident from our data. However, multidimensional analysis (Fig. 5) identified clearly different patterns in control and ALS (undSOD1 and mutSOD1) lymphoblasts, mainly involving parameters related to total SOD activity, intracellular oxidative stress (DCF fluorescence), catalase transcript levels, SOD1 protein levels (raw data), Nrf2 transcript levels, t-BHP (5000 μM)-induced loss of metabolic activity, and GR activity. It was possible to clearly separate ALS from control samples with these features. The two types of ALS samples (undSOD1 and mutSOD1) were more similar, although a perfect distinction was also achieved, both in the clustering (Fig. 6 b), and in the PCA (Fig. 6c) analyses, providing a perfect separation of samples belonging to each experimental group. This indicates that studying lymphoblast redox profiles may be helpful for patient stratification and precision medicine.

## 5 Conclusions

The results from this study revealed oxidative stress parameters in lymphoblasts from ALS patients that can be promising biomarkers for disease stratification or further exploited as therapeutic targets for personalized ALS drug development.

## Acknowledgements

This work was financed by the European Regional Development Fund (ERDF), through the COMPETE 2020 - Operational Programme for Competitiveness and Internationalization and Portuguese national funds via FCT – Fundação para a Ciência e a Tecnologia, under projects, PTDC/MED-FAR/29391/2017, POCI-01-0145-FEDER-029391, PTDC/BTM-SAL/29297/2017, POCI-01-0145-FEDER-029297, PTDC/BTM-ORG/0055/2021, DL57/2016/CP1448/CT0016 [TCO], UIDP/04539/2020, UIDB/04539/2020 and LA/P/0058/2020.

## Data availability

The datasets generated in this work are available at the figshare repository (https://figshare.com/projects/Oxidative_stress_profiling_of_ALS_lymphoblasts/132272)

## Abbreviations

ALS: Amyotrophic Lateral Sclerosis
ARE: Antioxidant Response Element
BCA: Bicinchoninic Acid
CSF: Cerebrospinal Fluid
DTT: Dithiothreitol
ECL: Enhanced Chemiluminescence reagent
EDTA: Ethylenediaminetetraacetic Acid
EGTA: Ethylene Glycol-bis (2-aminoethylether) N, N,’N, ’N-tetraacetic Acid
fALS: Familial ALS
GPx: Glutathione Peroxidase
GR: Glutathione Reductase
GSH: Reduced Glutathione
GSSG: Oxidized Glutathione
H_2_DCFDA: 2’,7’-Dichlorodihydrofluorescein Diacetate
HPLC: High-Performance Liquid Chromatograph
HRP: Horseradish Peroxidase
MDA: Malondialdehyde
mutSOD1: Mutant SOD1
NAM: Nicotinamide
Nrf2: Nuclear Factor Erythroid-2-related Factor
PBMCs: Peripheral Blood Mononuclear Cells
PBS: Phosphate-Buffered Saline
PCA: Principal Component Analysis
PMSF: Phenylmethylsulfonyl Fluoride
PVDF: Polyvinylidene Difluoride
sALS: Sporadic ALS
SDS: Sodium Dodecyl Sulfate
SOD1: Superoxide Dismutase 1
Ta: Annealing Temperature
TBA: Thiobarbituric Acid
t-BHP: tert-Butyl Hydroperoxide
undSOD1: Undetermined SOD1 Mutations
vit E: Vitamin E

## Statements & Declarations

### Author contributions

Filomena S.G Silva, Daniela Franco Silva, Ricardo Marques, Luis Segura and Inês Baldeiras performed experiments. Filomena S. G. Silva, Teresa Cunha-Oliveira, Tatiana Rosenstock, and Paulo J. Oliveira designed research and acquired funding. Filomena S. G. Silva and Teresa Cunha-Oliveira analyzed data. Filomena S. G. Silva and Teresa Cunha-Oliveira wrote the first draft of the paper. Teresa Cunha-Oliveira prepared the figures. All authors read and approved the final manuscript.

### Ethics approval and consent to participate

Not applicable.

### Consent to participate

Not applicable.

### Consent to publish

Not applicable.

### Availability of data and material

The full datasets obtained during the current study are available from the figshare repository (https://figshare.com/projects/Oxidative_stress_profiling_of_ALS_lymphoblasts/132272).

## Tables

**Supplemental Table 1:** List of cell samples and known characteristics.

| Sample | Control_C1 | undSOD1_C1 | mutSOD1_C1 | Control_C2 | undSOD1_C2 | mutSOD1_C2 | Control_C3 | undSOD1_C3 | mutSOD1_C3 |
| --- | --- | --- | --- | --- | --- | --- | --- | --- | --- |
| Coriell Reference RRID | ND14405<br>CVCL_GL09 | ND16061<br>CVCL_BS65 | ND13750<br>CVCL_BP71 | ND05102<br>CVCL_IB49 | ND12215<br>CVCL_BJ76 | ND10941<br>CVCL_BF63 | ND07879<br>CVCL_IC97 | ND11570<br>CVCL_BH13 | ND13750<br>CVCL_BP71 |
| Sex | F | F | F | M | M | M | M | M | M |
| Age | 46 | 46 | 46 | 46 | 46 | 46 | 26 | 27 | 26 |
| Description | Control | Patient with ALS | Patient with ALS | Control | Patient with ALS | Patient with ALS | Control | Patient with ALS | Patient with ALS |
| Identified mutations | ND | ND | mutSOD1: ALA4VAL | ND | ND | mutSOD1: ILE113THR | ND | ND | mutSOD1: ALA4VAL |

Available at https://doi.org/10.6084/m9.figshare.19146548

**Supplemental Table 2:**
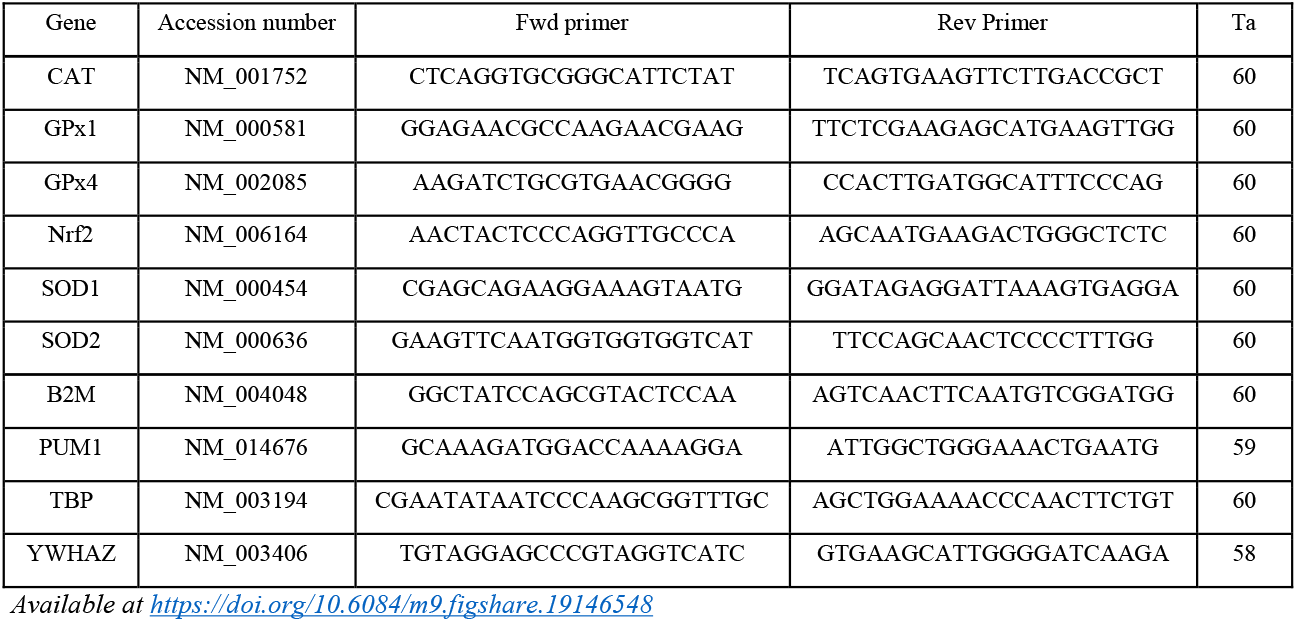
List of primers used in this work.

